# The cytochrome P450 enzyme WsCYP71B35 from *Withania somnifera* has a role in withanolides biosynthesis and defense against bacteria

**DOI:** 10.1101/2023.10.27.564345

**Authors:** H.B. Shilpashree, Ananth Krishna Narayanan, Sarma Rajeev Kumar, Vitthal Barvkar, Dinesh A. Nagegowda

**Author notes:** Address for correspondence: Dr. Dinesh A. Nagegowda.

## Abstract

The medicinal properties of Ashwagandha (*Withania somnifera* L. Dunal) are attributed to withanolides, which belong to the triterpenoid steroidal lactones class of compounds. Though it is proposed that intermediates of the universal phytosterol pathway are utilized by cytochrome P450 (CYP450) enzymes to form withanolides, studies on functional characterization of these enzymes has been sparse. This study reports the functional characterization of a CYP450 candidate from *W. somnifera* (WsCYP71B35) that exhibited induced expression in response to methyl jasmonate treatment and showed higher expression in tissues that accumulate withanolides. Biochemical assay with yeast microsomal fraction expressing recombinant WsCYP71B35 indicated no activity when phytosterols and their intermediate 24-methylene cholesterol were used as substrates. However, WsCYP71B35 catalyzed product formation with withaferin A, withanolide A, withanolide B, and withanoside IV among the tested substrates. Moreover, virus-induced gene silencing (VIGS) and transient overexpression of *WsCYP71B35* in *W. somnifera* leaves modulated the levels of withaferin A, withanolide A, and withanolide B, indicating the role of *WsCYP71B35* in withanolides pathway. Furthermore, VIGS of *WsCYP71B35* in *W. somnifera* reduced its tolerance to *Pseudomonas syringae* (DC3000) infection, whereas overexpression enhanced the tolerance to the bacterium in *W. somnifera* and transgenic tobacco. Overall, these results provide insights into the role of *W. somnifera* WsCYP71B35 in withanolides biosynthesis and bacterial defense.

## Introduction

Specialized terpenoids represent the largest and most diverse group of plant specialized metabolites and are often specific or unique to individual plant species or groups of species. They provide overall fitness to the plant in terms of their ecological roles and defense against stresses. Among the terpenoids, triterpenoids/steroids are the most numerous groups of plant metabolites and their structural diversity lies in the modification of side chains (Thimmappa et al. 2014). Besides their beneficial role to plants, these specialized triterpenoids have immense value to humans due to their medicinal properties (Banerjee et al. 2019). Withanolides are a group of secondary metabolites belonging to the class of tetracyclic triterpenoid steroidal lactones that contains more than 600 structurally distinct compounds, found mostly in plants of the Solanaceae family (Knoch et al. 2018). The best-known plant to produce withanolides is *Withania somnifera*, also known as Indian ginseng or Ashwagandha. It is a medicinal plant that has been used for over 3,000 years in traditional Indian medicine or Ayurveda and is also relevant in modern medicine (Bharti et al. 2021). Traditional healers have been using this plant for over 6000 years to treat a wide variety of ailments including asthma, fever, inflammation, abdominal pain, constipation, helminths, tuberculosis, cardiac disorders and to improve overall fitness (Joshi and Joshi 2021). In *W. somnifera,* withanolides are mainly found in leaves and roots ranging from 0.001 to 0.5 % of dry weights. Modern research has shown that a number of withanolides have promising pharmacological properties for the treatment of inflammation-associated chronic diseases including cancer, arthritis, autoimmune and neurodegenerative diseases (White et al. 2016; Wang 2020). So far, over 40 withanolides have been reported from this plant of which withaferin A and withanolide D have been reported to be the main active compounds contributing to the medicinal properties of Ashwagandha (Dhar et al. 2015). Both Withaferin A and Withanolide D have been reported to inhibit angiogenesis, Notch-1 and NFjB in cancer cells, and induce apoptosis in breast cancer cells (Kaileh et al. 2007; Koduru et al. 2010; Hahm et al. 2011). Plant extracts of *W. somnifera* as well as purified withanolides have demonstrated diverse pharmacological activities such as anti-inflammatory, anti-tumor, cardioprotective, neuroprotective and anti-bacterial properties (Mirjalili et al. 2009; Sehgal et al. 2012).

Despite their immense pharmacological and therapeutic potential, commercial exploitation of withanolides has been limited owing to low *in planta* accumulation (in the range of 0.001% to 0.5% of DW) resulting in limited availability in purified forms (Agarwal et al. 2018). Also, the accumulation of withanolides is influenced by various factors such as growth rate, tissue type, geographical and environmental conditions, and chemotype (Dhar et al. 2013). Moreover, successful metabolic engineering to improve withanolides production requires a complete understanding of the genes involved in the biosynthetic pathway. Structurally, withanolides are C_28_ steroids with ergostane skeleton in which C_26_ and C_22_, or C_26_ and C_23_ are oxidized in order to form a δ- or γ-lactone (Chen et al. 2011). Withanolides have been reported to be derived through the universal phytosterol pathway (Figure 1). It is proposed that the intermediates of phytosterol pathway derived via cycloartenol undergo various biochemical transformations such as hydroxylation and glycosylation postulated to be carried out by cytochrome P450 enzymes (CYP450s) and glycosyltransferases (GTs) leading to the biosynthesis of withanolides and withanosides, respectively (Lockley et al. 1976; Sangwan et al. 2008; Singh et al. 2015). CYP450 enzymes are a large class of heme-thiolate proteins that are ubiquitous in their presence across all genera of organisms and catalyze diverse reactions in pivotal molecular pathways. They have been studied and classified by Nelson, DR (2009) and available on https://drnelson.uthsc.edu/CytochromeP450.html and an estimated 80 CYP450s have been assigned biochemical functions related to the plant triterpene metabolism (Ghosh 2017).

**Figure 1.**
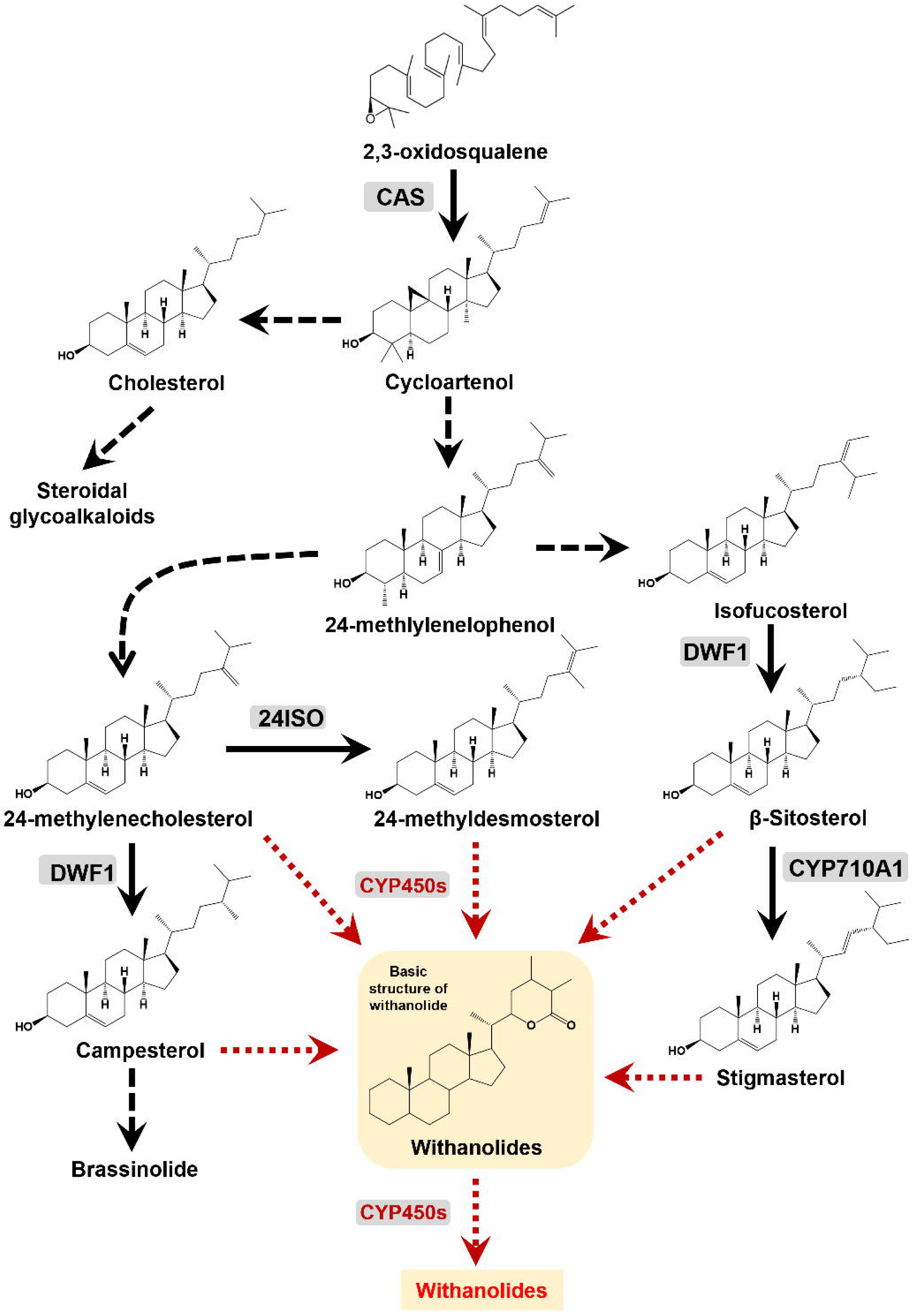
Overview of withanolides biosynthesis via phytosterol pathway in *W. somnifera*. Solid black arrows indicate single-step reactions, dashed arrows denote several steps, and dotted red arrows represent unidentified steps performed possibly by CYP450 enzymes. CAS, cycloartenol synthase; CYP710A1, C-22 sterol desaturase; DWF1, D24 sterol reductase; 24ISO, sterol Δ24-isomerase.

Efforts have been made by different research groups to identify and characterize CYP450s in *Withania somnifera.* Elicitor responsive *WsCYP98A* and *WsCYP76A* have been found to express abundantly in stalk and root, respectively, correlating in a positive manner to withanolides accumulation (Rana et al. 2014). In another study, it was shown that WSCYP93Id protein catalyzed the conversion of withaferin A to an unidentified hydroxylated product (Srivastava et al. 2015). Functional characterization of *WsCYP85A69* through miRNA and transient overexpression resulted in modulation of castasterone, stigmasterol and withanolides. Further, it was shown that WsCYP85A69 catalyzed *in vitro* conversion of 6-deoxocastasterone into castasterone (Sharma et al. 2019). Despite these studies, the knowledge on the biochemical and *in-planta* role of CYP450s in withanolides biosynthesis is poorly understood. Recently, we have performed molecular and *in planta* characterization of three CYP450 genes (*WsCYP749B1*, *WsCYP76* and *WsCYP71B10*) that showed their involvement in withanolides formation and defense in *W. somnifera* (Shilpashree et al. 2022). Here, we present the functional characterization of another CYP450 (WsCYP71B35) from *W. somnifera* and show its participation in withanolide biosynthesis and defense using biochemical, *in planta* silencing, and overexpression studies.

## Materials and Methods

### Plant material and methyl jasmonate treatment

Ashwagandha (*W. somnifera* cv. Poshita) (National Gene Bank, CSIR-CIMAP, India) seeds were sown by casting method in a seed bed tray. Planting mixture was composed of sterile soilrite, vermicompost and soil in the ratio of 1:1:1. Two-leaf staged seedlings were transplanted into pots and were maintained in growth chamber at 25°C with 16 h light and 8 h dark cycle and 70% humidity. Plants of appropriate leaf stage were used to carry-out VIGS and overexpression experiments. Methyl jasmonate (MeJA) treatment was performed using 10-week-old plants as described previously (Singh et al. 2017). Samples were harvested at different time (h) intervals and stored at 80°C until further use. To analyze the gene expression in different tissues, berries, flowers, leaves, roots and stems were collected from *W. somnifera* plant and stored in −80 °C for further use.

### Phylogenetic analysis

Amino acid sequences of cytochrome P450s belonging to CYP71 family that have been shown to be involved in secondary metabolism were used to construct phylogenetic tree. Neighborhood joining method was used for tree construction using MEGAX with default setting and bootstrap value changed to 1000 (Tamura et al. 2013). GenBank accession numbers for all CYP450 sequences used in phylogenetic tree are given in supporting information (Table S1).

### RNA isolation, cDNA synthesis and qRT-PCR analysis

Total RNA from different tissues was extracted from 100 mg tissue using the TRIzol reagent or SpectrumTM Plant Total RNA kit (Sigma-Aldrich, United States) following manufacturer’s instructions. On-column DNase digestion was performed with DNase I (Sigma-Aldrich, St. Louis, MO, United States) to remove contaminating genomic DNA. cDNA was synthesized with 2 µg total RNA and random hexamer primers using High Capacity cDNA Reverse Transcription kit (Applied Biosystems, United States) as per manufacturer’s instructions. A linear range of cDNA was determined with *18S* rRNA (endogenous control) and used for qRT-PCR analysis. qRT-PCR analysis was carried out with gene specific primers that were designed outside the gene region used for cloning into pTRV2 vector (Table S2). The reaction was performed using 2X SYBR green mix (Thermo Scientific, USA) and run in StepOne Real-Time PCR System (Applied Biosystems, USA). The qRT-PCR conditions used were as follows: 94°C for 10 min, followed by 40 cycles of 94°C for 15s and 60°C for 45 sec. Fold-change differences in gene expression were analyzed using the comparative cycle threshold (*C*_t_) method.

### Generation of VIGS and overexpression constructs

For generation of VIGS construct, a 500 bp fragment corresponding to WsCYP71B35 was amplified from leaf cDNA using gene specific primers. The amplicon was cloned into pGEMT-easy vector, sequence confirmed by nucleotide sequencing, and subsequently sub-cloned into *Xba*I and *Xho*I sites of pTRV2 plasmid that was procured from TAIR (www.arabidopsis.org), resulting in pTRV2::WsCYP71B35 construct (Figure S1, Table S2). To generate the plant overexpression construct, first the open reading frame (ORF) of WsCYP71B35 was PCR amplified using leaf cDNA, and the resulting amplicon was cloned into pJET1.2/vector and confirmed by nucleotide sequencing. The WsCYP71B35 fragment was restriction digested from pJET1.2 construct using *Xba*I and *Sac*I and subcloned into the respective sites of pBI121 vector under 35S promoter to form pBI121::WsCYP71B35 construct (Figure S1, Table S2). The plasmids (pTRV1, pTRV2, and pBI121) and constructs (pTRV2::WsCYP71B35 and pBI121::WsCYP71B35) were individually transformed into *A. tumefaciens* by freeze thaw method.

### VIGS and transient overexpression of WsCYP71B35

VIGS and transient overexpression of WsCYP71B35 was performed as described earlier (Singh et al. 2017). For VIGS, the transformed *A. tumefaciens* (strain GV3101) cultures were grown in 100 mL YEP medium at 28°C. The overnight grown cultures were harvested by centrifugation and resuspended in infiltration buffer (10 mM MgCl_2_, 10 mM MES and 100 µM acetosyringone, pH 5.6) to a final OD600 of 1.6 and incubated at 28◦C for additional 4-5 h. Infiltration was performed using 1:1 ratio of *Agrobacteria* cultures harboring TRV1 and TRV2 or its derivatives using 1-mL needleless syringe on the abaxial side of leaves in 4-leaf-staged plants. Leaves exhibiting viral infection phenotype were harvested 30 days post infiltration, and used for transcript and metabolite analyses. Transient overexpression was performed by co-infiltration of *Agrobacteria* harboring pBI121::*WsCYP71B35* and p19, and pBI121 and p19 vectors (control). Briefly, overnight Agrobacteria cultures were pelleted and resuspended in infiltration buffer (50 mM MES, 2 mM Na3PO4, 0.5% glucose, and 100 µM acetosyringone with pH adjusted to 5.6) to a final OD600 of 0.2 and incubated at 28°C for 3-4 hour. Leaf infiltration of Agro-suspension was performed as described above into the first pair of leaves of 6-8 leaf-staged plant and placed in dark condition for 48 h. The infiltrated leaves were harvested and stored in −80°C for metabolite and transcript analyses (Singh et al. 2015).

### Overexpression of WsCYP71B35 in yeast and CYP450 assay

The open reading frame of WsCYP71B35 was amplified and cloned into *Bam*HI and *Eco*RI sites of yeast expression vector pYEDP60u resulting in pYEDP60u::*WsCYP71B35* (Figure S1, Table S2). The empty vector and the overexpression constructs were transformed into *Saccharomyces cerevisiae* WAT11 strain (with yeast reductase replaced by the ATR1 reductase from Arabidopsis thaliana, under GAL10-CYC1 promoter andCPR1 terminator)(Urban et al. 1997). The confirmed colonies were cultured using high-density procedure as described (Pompon et al. 1996). Briefly, an individual colony was streaked onto a SG(A)I plate. A loopful of culture from the plate was inoculated into 30 ml of SG(A)I and grown to stationary phase (overnight). A 1:50 dilution was made into 500 ml YPGE medium (yeast extract-10 g, Bacto-Peptone- 10 g, glycerol- 20 mL and ethanol- 10 mL per liter of H2O) and cells were grown at 30°C in a shaking incubator until cell density (OD600) reached 8 x 107 cells per ml (∼24-36 h). Induction was initiated by the addition of 10% (v/v) of sterile aqueous solution of galactose (20 mg/mL). The induction was continued for 8-15 h until the cell density reached 2-5 x 108 cells per ml. The cultures were pelleted and cells were mechanically broken using centrifugation for microsomal isolation. Protein content in isolated microsomes was determined by Bradford’s method using Bovine Serum Albumin (BSA) as a standard. CYP450 assay was performed using potassium phosphate buffer (50 mM, pH 7.4) containing: 1 mM NADPH, and 100 uM of potential substrate (phytosterol intermediates and different commercially available withanolides/withanosides). The reaction was initiated by addition of 0.2-0.5 mg of microsomal fraction of *S. cerevisiae* harboring either pYEDP60u (control) or pYEDP60u::*WsCYP71B35*, and the mixture was incubated at 30°C for 1 h in a water bath. The reaction was terminated by chilling the mixture on ice and adding 3N HCl. The reaction products were extracted by the addition of chloroform: methanol (7:3) and the extract was used for detection using HPLC analysis as described below.

### Extraction and analysis of withanolides

Withanolides extraction and analysis from *W. somnifera* was performed following Singh et al. (2015). Leaf tissues were oven dried at 55°C and 20 mg of dried tissue was ground using 1 mL absolute methanol, sonicated and the supernatant was collected. The ground tissue was re-extracted twice and obtained methanolic extracts were pooled in a scintillation vial and allowed to evaporate. The remaining residue was resuspended in 3 mL of 70% methanol. This was extracted thrice with 3 mL chloroform. The lower layer of chloroform extract (9 mL) was decolorized using charcoal and dried. To the dried samples, 200 µL of 7:3 chloroform:methanol was added and used for HPLC analysis. Whereas, the CYP450 enzyme assay products were directly used for HPLC analysis. Withanolides were analyzed using HPLC (Model: Nexera, Shimadzu, Kyoto, Japan) fitted with C-18 column (250 mm x 4.60 mm, 5 µm) as per the program described previously. Briefly, Potassium dihydrogen phosphate buffer (1.02 mM) and acetonitrile were used as aqueous and organic solvents. Gradient program was used to separate withanolides. Initially, solvent A concentration was set at 95%, then changed to 55%- 20% at 18 and 25 min. 20% of solvent A was maintained for the next 10 min, later it was increased to 55% at 35th min and 95% at 40th min. The flow rate was set to 1.5 mL/min with UV detection at 227 nm. All standards of withanolides were from Natural Remedies Pvt. Ltd. (Bangalore, India) and the standard stock concentration was 1 mg/mL. To 36 µL of withanolides extract, 4 µL of 1 mg/mL catharanthine was added and 20 µL of this mixture was injected into the HPLC system. The area of individual withanolides was determined after normalizing with the peak area of the internal standard, catharanthine (Sigma-Aldrich) (Singh et al. 2015, 2017).

### Bacterial growth curve assay

Bacterial growth assay was performed using the model plant pathogen *P. syringae* pv. tomato DC3000. The bacterium was cultured in 5 mL nutrient broth and grown in an incubator shaker at 28°C. The overnight grown culture was centrifuged and the pellet was resuspended in 10 mM MgCl2 to achieve an OD_600_=0.002. Bacterial infiltration and cfu determination were performed as described previously (Singh et al. 2015, 2017). Leaf discs from infected and buffer infiltrated control plants were collected 3 days post infiltration and used for assessing the bacterial infection rate. Leaf discs were homogenized using 10 mM MgCl2 and the cfu/cm2 was determined by plating serial dilutions of leaf extracts on Pseudomonas-specific agar plates.

### Generation of transgenic tobacco

Leaf disc co-cultivation with A. tumefaciens harboring pBI121 and pBI121::WsCYP71B35 was carried out to transform tobacco as described in Singh et al., 2017. Transformed shoots regenerated on MS (Murashige and Skoog 1962) medium supplemented with kanamycin (100 mg/L-1) selection media were transferred to ½ strength MS agar for rooting. Plantlets having well-developed roots were removed from the bottle and transferred to pots containing sterile soilrite mix for hardening. Further, the plants were PCR screened and positive lines were transferred to growth chamber. Seeds were collected from T0 lines and used to generate T1 plants. PCR-screened positive lines of T1 plants were used for transcript and bacterial growth analyses (Singh et al. 2017).

### Statistical analysis

Mean, standard deviation and standard error was calculated using GraphPad QUICKCALC online tool. The statistical significance between control and treated samples was calculated using unpaired Students t- test. Asterisks indicate significant difference between samples (\**P*<0.05; \*\**P*<0.01; and \*\*\**P*<0.001).

## Results

### Identification and sequence analysis WsCYP71B35

In our previous work, three WsCYP450 genes having induced expression were identified and characterized at *in planta* level (Shilpashree et al. 2022). Here, a fourth CYP450 candidate belonging to CYP71 family (*WsCYP71B35*) is functionally characterized. *WsCYP71B35* contains an open reading frame of 1515 bp that encodes a protein of 504 amino acids with a calculated molecular mass of 57190.50Da (GenBank accession number: MW298520.1). Analysis of the deduced amino acid sequences of the encoded protein showed the presence of [FW]-[SGNH]-x- [GD]-{F}-[RKHPT]-{P}-C-[LIVMFAP]-[GAD] consensus pattern for cysteine heme-iron ligand signature, where C is the heme iron ligand (Figure 2). Phylogenetic analysis with candidates of CYP71 family from other plant species indicated that WsCYP71B35 has 58.57% identity with *Hyoscyamus muticus* CYP71 followed by 46.8% identity with MgCYP71D95 from *Mentha x gracilis* (Figure 2A). WsCYP71B35 possessed the most conserved motif among CYP450 enzymes, which is the heme-binding region FxxGxRxCxG (also known as the CxG motif), containing the axial Cys ligand that binds to the heme (Figure 2B). Also, WsCYP71B35 contained ExxR and PER motifs that form the E-R-R triad, which is important for locking the structure of the heme pocket in place and ensuring the stabilization of the core structure (Córdova et al. 2017). Prediction of subcellular localization using different tools indicated that *CYP71B35* is membrane targeted, which is a characteristic of CYP450 (Table S3).

**Figure 2.**
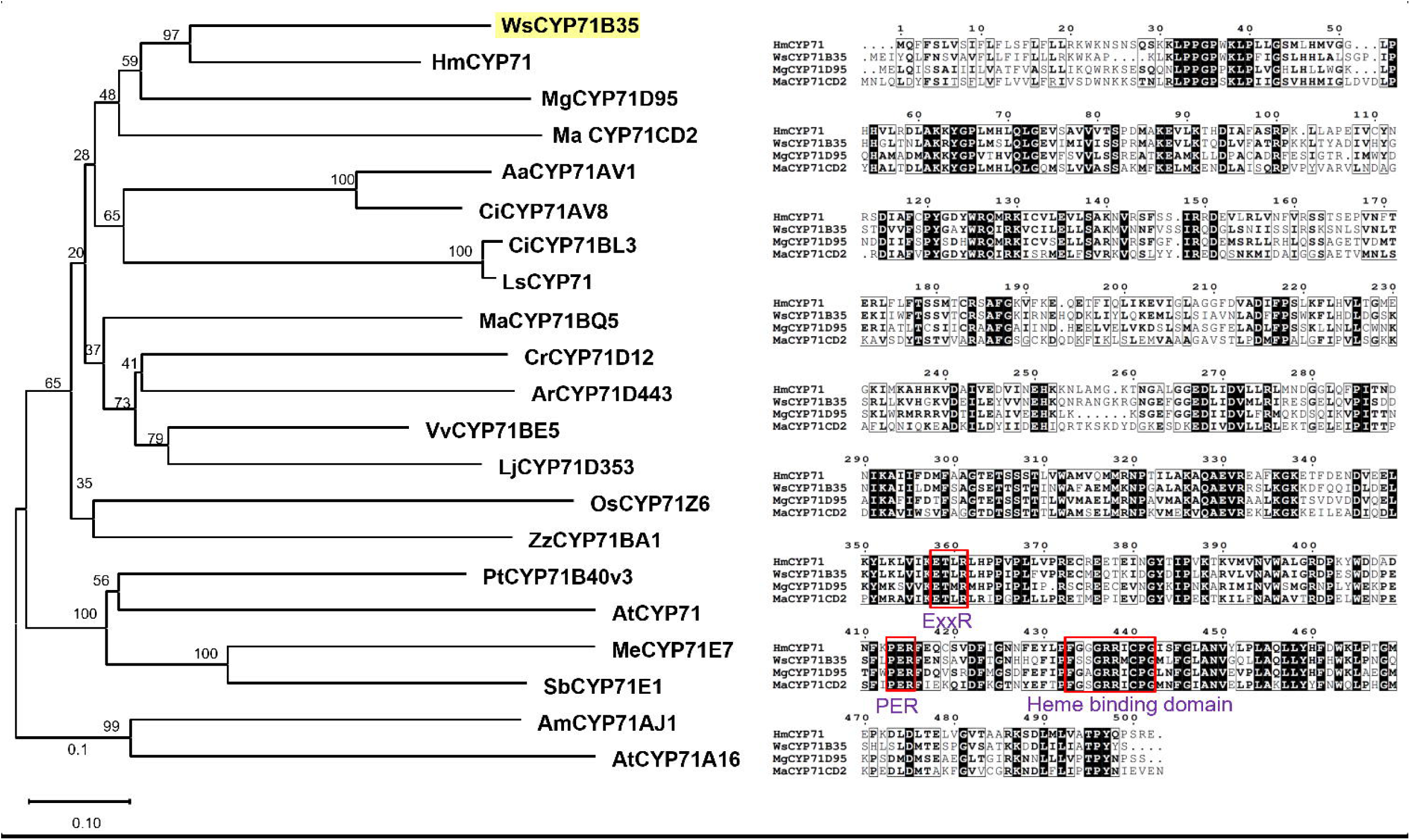
Sequence analysis of WsCYP71B35 with other related plant CYP450 enzymes. (A) Phylogenetic tree showing the relationship of *WsCYP71B35* with other plant CYP450s of the CYP71 family. The tree was constructed using MEGAX software (Kumar et al., 2016), and statistical reliability of individual nodes of the tree was assessed by bootstrap analyses with 1000 replicates. The GenBank accession numbers of sequences used for constructing the phylogenetic tree are shown in Table S1. (B) Multiple sequence alignment of *WsCYP71B35* with related CYP71 family enzymes from *Catharanthus roseus* (ACM92061)*, Hyoscyamus muticus* (ABS00393), and *Manihot esculenta*, (Q6XQ14). The characteristic heme-biding domain of CYP450s is shown in a red box.

### Expression of WsCYP71B35 in response to MeJA and in different tissues

MeJA is a known inducer of pathway genes and transcription factors involved in secondary metabolism including genes related to withanolides biosynthesis. To determine whether expression of *WsCYP71B35* is induced in response MeJA qRT-PCR was performed using leaf tissues collected at different time intervals after MeJA treatment. It was found that *WsCYP71B35* exhibited a drastic induced expression of almost 80-fold at 6 h upon exposure to MeJA (Figure 3A). However, *WsCYP71B35* expression was reverted back to 0 h level at 12 h, and remained the same at 24 h and 48 h after MeJA treatment. The observed MeJA-induced expression of *WsCYP71B35* indicated its possible involvement in secondary metabolism and possibly in withanolides biosynthesis (Figure 1). Further, expression analysis using different tissues revealed that *WsCYP71B35* possessed highest expression in tissues that predominantly accumulate withanolides, i.e., leaf (∼70-fold), followed by fruit (∼50 fold), flower and root (∼30-fold each) with respect to the stem where the least expression was seen (Figure 3B).

**Figure 3.**
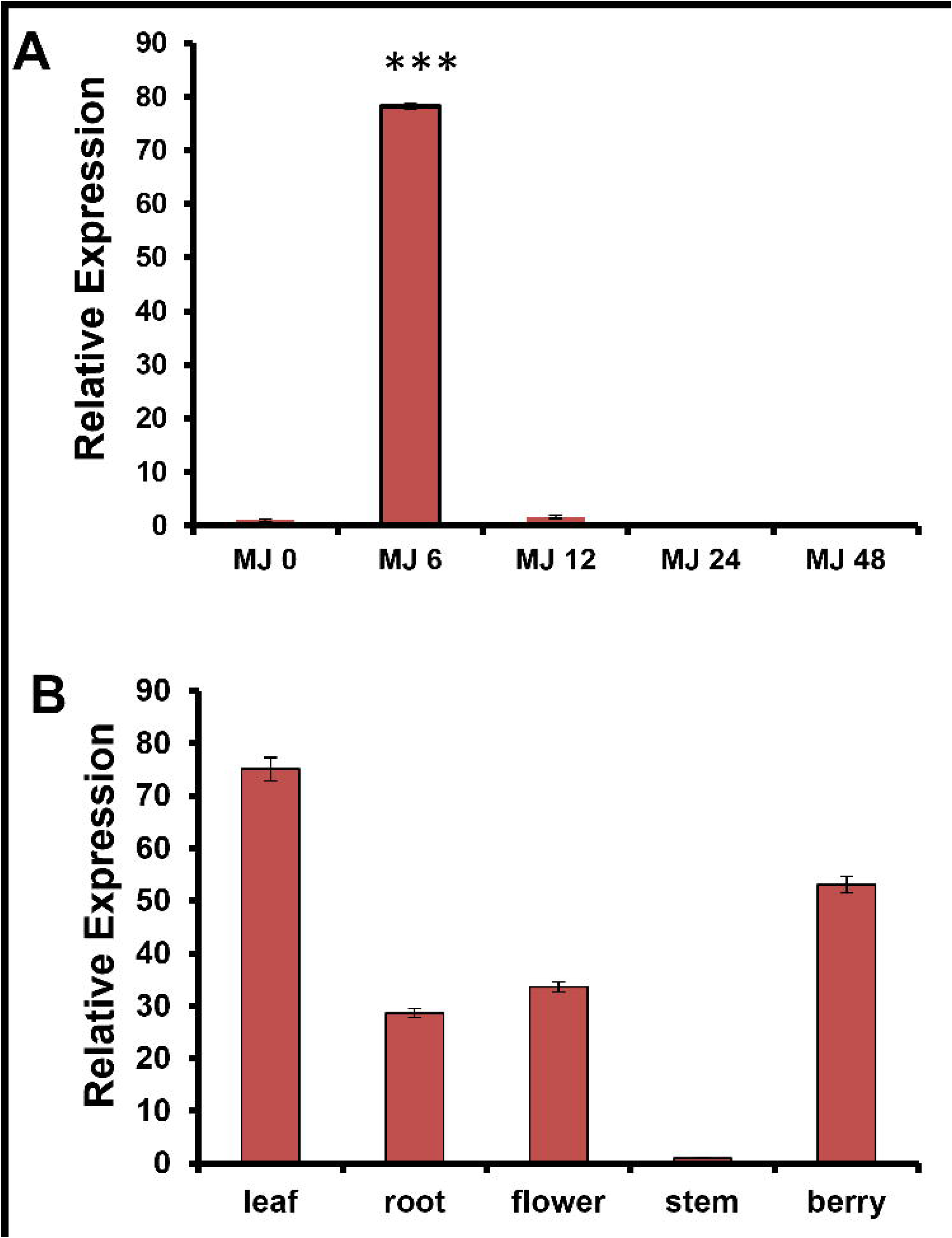
Gene expression analysis of *WsCYP71B35*. (A) Effect of methyl jasmonate treatment on *WsCYP71B35 mRNA* levels at time intervals of 0, 6, 12, 24, and 48 hours. The expression is represented relative to the control leaf tissue. (B) Relative transcript abundance of *WsCYP71B35* in different tissues of *W. somnifera*. In each graph, the tissue having the least *Ct* was set to one-fold to determine the relative abundance of transcripts in other tissue. *18S rRNA* was used as an internal reference for normalization. The data shown are from three independent experiments. Student’s *t*-test: *, *p*<0.05; **, *p*<0.01. Error bars indicate mean ± SE.

### Biochemical characterization of WsCYP71B35

To understand the biochemical function of WsCYP71B35 enzyme, the ORF was cloned into yeast overexpression vector and the recombinant protein was expressed in yeast. Since withanolides are reported to be derived from the universal sterol pathway in *W. somnifera*, phytosterols such as campesterol, cholesterol, stigmasterol, beta-sitosterol and one of their intermediates 24-methylenecholesterol (presumed to be the precursor of withanolides) were used as substrates. Enzyme assay using yeast microsomal fraction expressing recombinant WsCYP71B35, the phytosterol substrates, and NADPH as cofactor, followed by HPLC analysis of the reaction products indicated no activity with any of the tested phytosterols. Further, independent CYP450 assay with seven different withanolides (withanolide A, withanolide B, withanone, 12- deoxywithastramonolide, withanodise IV, withaferin A and withanoside V) as substrates and NADPH cofactor, showed that four substrates among the tested seven exhibited product formation. It was found that WsCYP71B35 catalyzed the product formation in the presence of withaferin A, withanolide A, withanolide B, and withnoside IV as substrates (Figure 4). Comparison of UV-spectra revealed that the products formed in each assay were different from each other (Figure S2).

**Figure 4.**
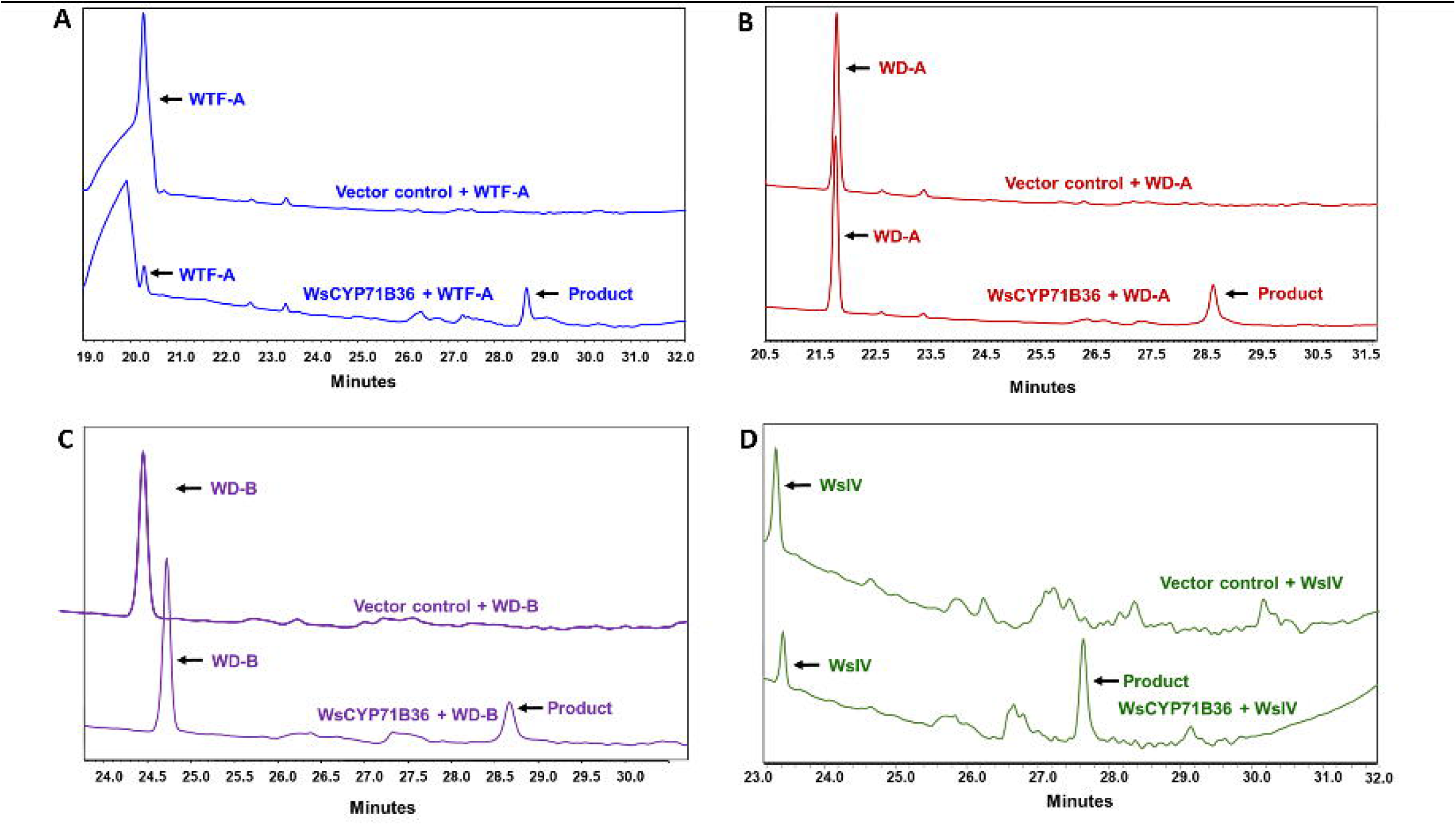
HPLC analysis of reaction products of CYP450 assay. The yeast microsomal fraction expressing the recombinant *WsCYP71B35* along with different withanolides, A) Withaferin A, B) Withanolide A, C) Withanolide B and D) Withanoside IV as substrates were used in the assay. Assay with substrates that showed product formation are only shown here. The chromatograms at the top and bottom of each sub-figure represent reactions using microsomal fraction of yeast transformed with empty vector (control) and pYeDP60u::*WsCYP71B35* construct, respectively.

### Effect of silencing and overexpression of WsCYP71B35 on withanolide biosynthesis

To determine the *in-planta* role of *WsCYP71B35* in withanolide biosynthesis, virus induced gene silencing (VIGS) was performed. As VIGS is prone to off target silencing (Dinesh-Kumar et al. 2003), the sequence region unique to *WsCYP71B35* was chosen to generate the silencing constructs. Thirty days post-infiltration, leaves of similar developmental stages exhibiting typical viral infection symptom were collected along with empty vector control for transcript and metabolite analyses. qRT-PCR analysis showed 80% reduction in the expression of *WsCYP71B35* compared to EV controls (Figure 5A). The degree of silencing of WsCYP450s in this study was comparable to the VIGS of other related genes of *W. somnifera* reported in earlier studies as well as our own previous study (Singh et al. 2015, 2017; Agarwal et al. 2018; Shilpashree et al. 2022). Subsequent analysis of metabolites through HPLC indicated that VIGS of *WsCYP71B35* did not significantly alter the levels of withanolides in the plants except for withanolide A. In the silenced plants, it was observed that withanolide A levels had a 75% reduction as compared to EV (Figure 5B). In addition, withaferin A showed an increased level that was not statistically significant in *WsCYP71B35-*vigs plants. In continuing to characterize the role of *WsCYP71B35 in planta*, we proceeded to overexpress this gene by agroinfiltration in *W. somnifera* leaves. Two days post infiltration, leaves were collected for metabolite and transcript analyses. qRT-PCR analysis showed that *WsCYP71B35* was overexpressed almost 8- fold in the infiltrated tissue (Figure 5C). HPLC analysis of the metabolites revealed an increased levels of withanolide A by around 60% and surprisingly, a decreased level of withanolide B by about 90% (Figure 5D).

**Figure 5.**
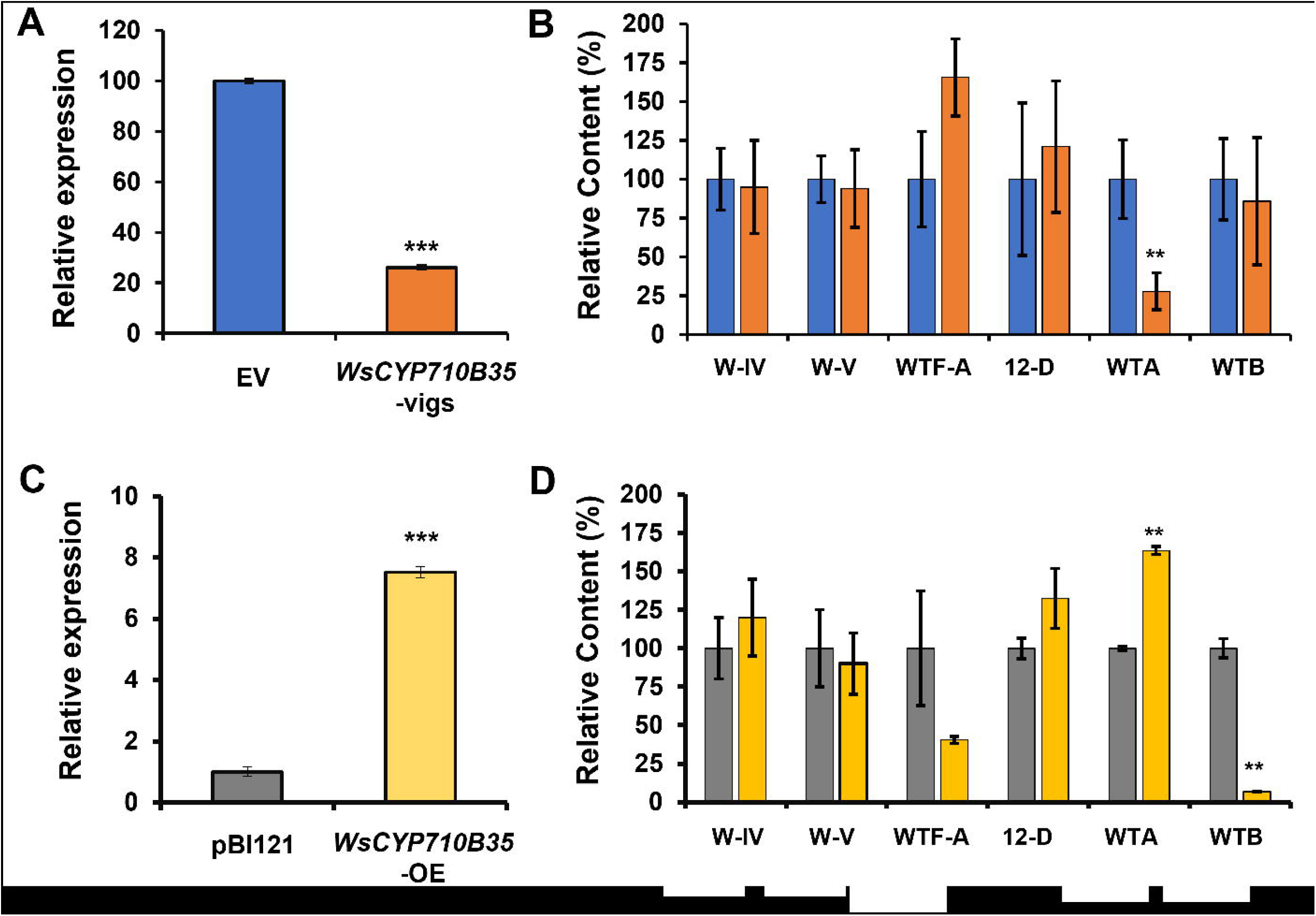
Effect of silencing and overexpression of *WsCYP71B35* on withanolide content. (A) Quantitative reverse transcription polymerase chain reaction (RT-qPCR) analysis of *WsCYP71B35* expression in silenced leaves. Expression level of *WsCYP71B35* was normalized to 18S rRNA and set to 1 in an empty vector (EV) control to determine the relative reduction in *WsCYP71B35*-silenced leaves. (B) High performance liquid chromatography (HPLC) quantification of different withanolides in control and *WsCYP71B35*-silenced leaves. (C) Quantitative reverse transcription polymerase chain reaction (RT-qPCR) analysis of *WsCYP71B35* expression in overexpressed (OE) leaves. (D) withanolides levels in control and *WsCYP71B35* overexpressing leaves quantified through HPLC. Levels of different withanolides are represented relative to EV. The peak area of individual withanolides was determined after normalizing with peak area of internal standard (catharanthine). Wit-IV, withanoside IV; Wit-V, withanoside V; Wit(de)-A, withanolide A; Wit(de)-B, withanolide B; Wit(in)-A, withaferin A; 12d-wit, 12-deoxywithastramonolide. The results shown are from (a) three, (b) five and (d) 10 independent experiments. Student’s t-test: *, *p*<0.05; ***, *p*<0.001. Error bars indicate mean SE

### Role of WsCYP71B35 in bacterial defense

In our previous studies in *W. somnifera*, it was observed that VIGS and overexpression of genes related to phytosterol and withanolides pathway modulate the expression of defense related genes and affect pathogen defense (Singh et al. 2015, 2017). Moreover, withanolides have been shown to possess antibacterial and antifungal properties (Choudhary et al., 1995). As silencing and transient overexpression of *WsCYP71B35* resulted in modulation of withanolides content, we proceeded to study their effect on *PR* genes. We focused on salicylic acid (SA)- dependent *PR1* and jasmonate-dependent *PR3* genes. It was observed that *PR1* and *PR3* expression did not show statistically significant changes in expression in the silencing (VIGS) background. In stark contrast, overexpression of *WsCYP71B35* led to significant upregulation of both *PR1* and *PR3* by 6.2 and 16.9-fold respectively (Figure 6). Following this, a bacterial growth assay was performed to determine *WsCYP71B35s*’ role in defense. Fully expanded leaves from *WsCYP71B35*-vigs and EV control plants were infiltrated with *P. syringae* DC3000. Leaves from silencing background developed severe disease symptoms and sustained more tissue damage than EV leaves. Further, bacterial growth assay using extract isolated from infiltrated leaves 3 dpi showed that *WsCYP71B35-*vigs samples exhibited drastic increase in the growth of *P. syringae* DC3000 than that of EV control. The log cfu/cm^2^ for *WsCYP71B35*-vigs was found to be 7.1 whereas it was ∼6 cfu/cm^2^ for EV control (Figure 7A). Leaves overexpressing *WsCYP71B35* remained healthy, and exhibited no necrotic phenotype compared to control at 4 dpi of *P. syringae* DC3000. Subsequent bacterial growth assay revealed a significant reduction in the growth of *P. syringae*. The log cfu/cm^2^ in *WsCYP71B35* overexpressing tissue was 5.3 while it was 7.2 in vector control (Figure 7B). The observed phenotype and bacterial growth in *P. syringae* infiltrated samples correlated with elevated and reduced expression of *WsCYP71B35* (Figure 5A and 5C).

**Figure 6:**
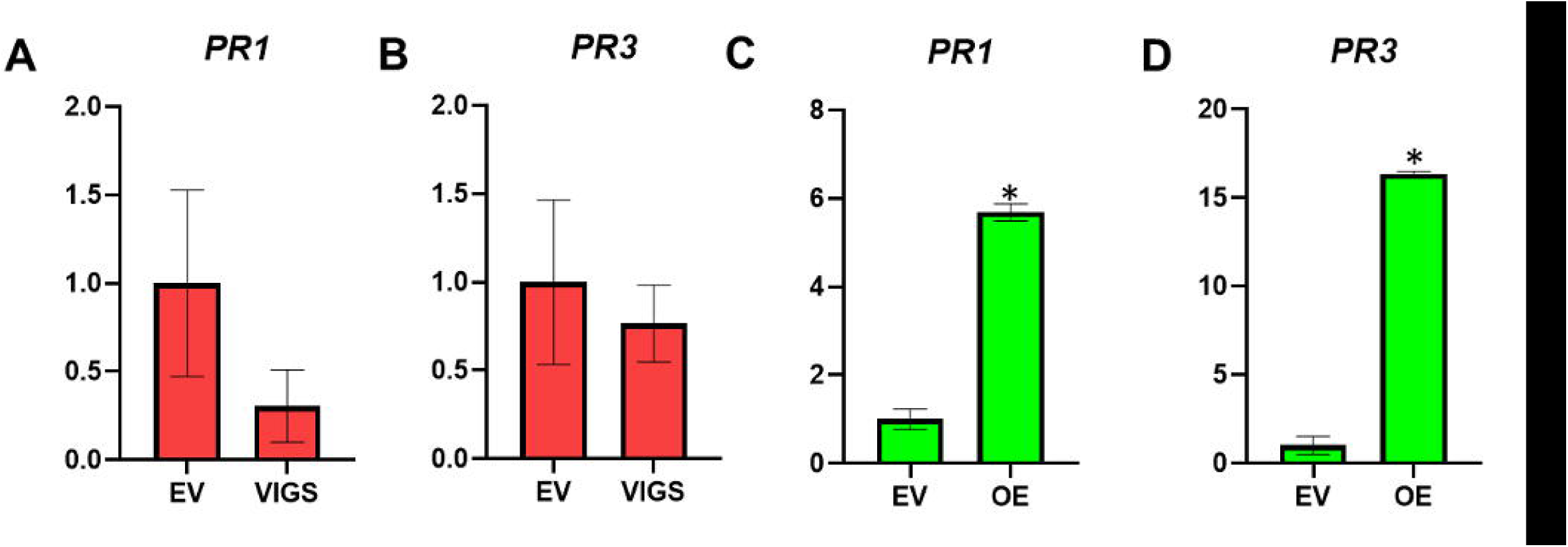
Relative expression of *PR* genes in *WsCYP71B35* modulated plants. A) Expression of PR1 and B) *PR3* in VIGS background. C) Expression of PR1 and D) *PR3* in overexpression background. Relative gene expression was analysed using qRT-PCR and calculated using the 2^-ΔΔCt^ method. Graphs were plotted using mean, error bars show□±□SE from three independent experiments. Statistical significance represented as \**p*<□0.01.

**Figure 7.**
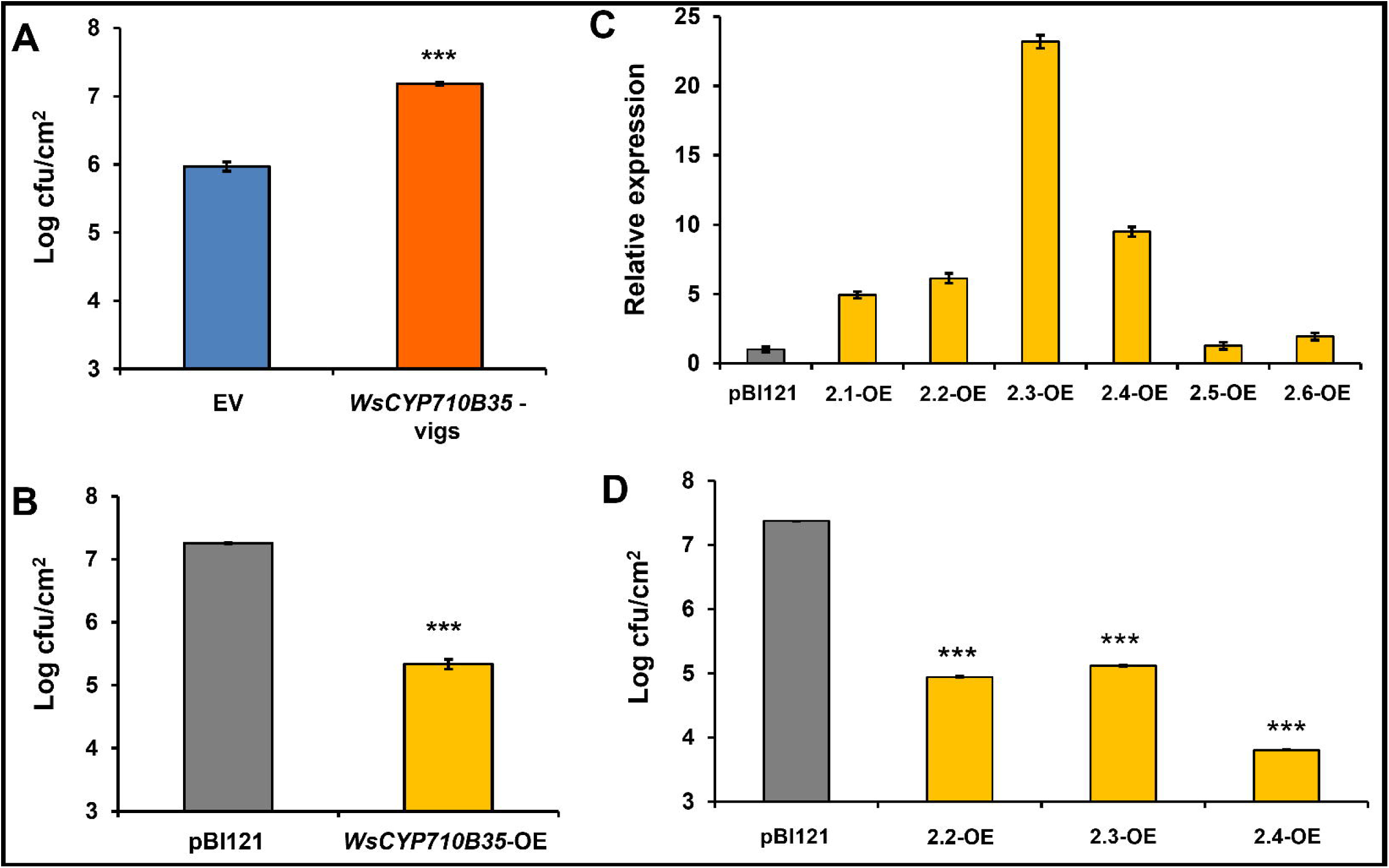
WsCYP71B35 confers tolerance to bacterial growth in *W. somnifera* and tobacco. *Pseudomonas syringae* growth assay in *WsCYP450s* silenced (A) and overexpressing (B) *W. somnifera* leaves. (C) Expression level of *WsCYP71B35* in different transgenic tobacco lines, normalized to internal reference *NtEF1*α and represented as expression relative to pBI121 control that was set to 1. (D) Bacterial growth assay of transgenic tobacco lines with *P. syrinage* (DC3000) strains. The bacterial growth (CFU) at 3 dpi was obtained by plating serial dilutions of leaf disc extracts. Error bars show mean□±□SE of three independent experiments. Statistical significance represented as \**p*<□0.05, \*\**p*<□0.01, \*\*\**p*<□0.001, a significant difference.

### Overexpression of WsCYP71B35 in transgenic tobacco confers tolerance to bacteria

Heterologous overexpression of plant CYP450s has been shown to confer tolerance to various stresses including tolerance against bacteria (Pandian et al. 2020). To further investigate the defensive role of *WsCYP450s* in heterologous system, independent transgenic tobacco lines overexpressing *WsCYP71B35* were generated. The transcript abundance of *WsCYP71B35* gene in 6 independent lines was determined by RT-qPCR and was found to be between 1- to 24-fold (Figure 7C). The three highest expressing lines and control plants were chosen for bacterial growth assays. Analysis of bacterial growth using extracts isolated from leaf discs of *P. syringae* infiltrated leaves at 3 dpi significantly repressed the multiplication of *P. syringae* as compared to that of empty vector infiltrated plants (Figure 7D). Similar to our results, overexpression of *Panax ginseng PgCYP76C9* conferred enhanced resistance to *P. syringae* in transgenic *Arabidopsis* (Balusamy et al. 2017).

## Discussion

Withanolides are bioactive constituents of *W. somnifera* belonging to triterpenoid steroidal lactone class of plant specialized metabolites. The terminal phytosterol pathway genes responsible for the biosynthesis of these immensely important molecules are yet to be identified. It is speculated that precursors derived from the universal sterol pathway undergo hydroxylation and other biochemical modifications postulated to be carried out by WsCYP450 enzymes leading to the formation of diverse withanolides (Srivastava et al. 2015). Therefore, understanding the specific WsCYP450s involved is essential to elucidate the biosynthesis of these valuable compounds. In this study, we set out to identify and characterize WsCYP450s involved in withanolides biosynthesis through biochemical, molecular and *in planta* studies. The newly identified WsCYP71B35 possessed the characteristic cysteine heme-iron ligand domain structure, and belonged to the CYP71 clan, which is the largest set of P450s in plants (Nelson et al., 2004). The families and subfamilies within the CYP71 clan have diverged remarkably during plant evolution (<u>Nelson et al., 2004</u>; <u>Nelson and Werck-Reichhart, 2011</u>), and are known to have diverse roles in the biosynthesis of terpenoids, phytoalexin, as well as indolic derivatives, cyanogenic glucosides, flavonoids, aldoximes and nitriles (Nelson and Werck-Reichhart 2011; Hamberger and Bak 2013; XU et al. 2015). Phylogenetic analysis showed that the closest related enzyme to WsCYP71B35 was premna-spirodiene oxygenase from *Hyoscyamus muticus* (HmCYP71) which shared 58.57% sequence identity. HmCYP71 was characterized to carry out the regio-specific (C-2) hydroxylation of several eremophilane-type (decalin ring system) sesquiterpenes including 5-*epi*-aristolochene (Takahashi et al. 2007). The second closest related enzyme in the tree was (-)-(4*S*)-limonene-3-hydroxylase from *Mentha x gracilis* (MgCYP71D95) which also carries out regio-specific hydroxylation of limonene at C-3 to (-)-*trans*-isopiperitenol (Bertea et al. 2003). Further, WsCYP71B35 also showed homology to the CYP71 enzymes involved in triterpenoid metabolism (Malhotra and Franke, 2022). They included *Arabidopsis* CYP71A16 that converts marneral/marnerol to 23-hydroxymarneral/23-hydroxymarnerol (Kranz-Finger et al. 2018), *Melia azedarach* CYP71BQ5 and CYP71CD2 which utilize dihydroniloticin and tirucalla-7,24-dien-3β-ol, respectively, forming corresponding products, melianol and dihydroniloticin (Hodgson et al. 2019), *Lotus japonicas* CYP71D353 that catalyzes the conversion of dihydrolupeol/20-hydroxylupeol to 20-hydroxylupeol/20-hydroxybetulinic acid (Krokida et al. 2013), and *Ajuga reptans* CYP71D443 that is involved in the conversion of 3β-hydroxy-5β-cholestan-6-one to 3β,22R-dihydroxy-5β-cholestan-6-one (Tsukagoshi et al. 2016). Given these similarities of WsCYP71B35 with other CYP71 enzymes involved in specialized terpenoid metabolism, it may be assumed that WsCYP71B35 may also possess regio-specific hydroxylation activity of withanolides.

Generally, expression of pathway genes involved in plant secondary metabolism and the corresponding secondary metabolites are induced in response to phytohormone elicitor MeJA. *W. somnifera* is no exception to this phenomenon wherein withanolides accumulation and expression of related genes have been shown to be induced in response to biotic and abiotic stresses (Dhar et al. 2015), and also in response to MeJA (Bhat et al. 2012; Dhar et al. 2014; Singh et al. 2015, 2017). Similarly, genes of phytosterol biosynthesis and a WRKY transcription factor were highly induced in response to MeJA (Singh et al. 2015, 2017). Also, genes encoding CYP450s that have been implicated in withanolides formation also are shown to be induced in response to MeJA treatment (Dhar et al. 2015). Similar to the above reports, the expression of *WsCYP71B35* was induced at 6^th^ hour of MeJA treatment and was higher in all tissues that are known to accumulate withanolides, indicating its possible role in withanolides biosynthesis (Figure 3). It has been reported that withanolides are formed via the universal phytosterol pathway, and 24-methylene cholesterol is suggested to be the starting precursor for all downstream withanolides (Lockley et al. 1976). Since CYP450 enzymes are known to utilize wide range of substrates, biochemical assay using yeast microsomal fraction expressing WsCYP71B35 with different phytosterols was performed. This revealed that there was no activity with any of the phytosterol substrates. The fact that this enzyme did not form any product when incubated with phytosterols indicated that they are likely to be involved in the formation of downstream and eventual withanolide products using the intermediate substrates made by other CYP450 enzymes of the metabolic pathway. However, when withanolides were employed as substrates, WsCYP71B35 exhibited the ability to catalyze the conversion of withaferin A, withanolide A, withanolide B, and withanoside IV, resulting in the formation of a distinct peak corresponding to an unidentified product, possibly resulting from hydroxylation (Figure 4). Elucidating the chemical structures of these products could shed light into the role of WsCYP71B35 in the withanolide biosynthetic pathway. However, as the withanolide biosynthesis is a complex pathway, deciphering the correct substrate for each CYP is difficult. For instance, a CYP450 from *W. somnifera* WSCYP93Id was found to convert withaferin A to a hydroxylated product, putatively thought to be 17-hydroxy withaferin A based on the correspondence of peak retention time of the product (Srivastava et al. 2015). In our attempts to identify the enzyme products of the biochemical assay using LC-ESI-MS/MS, we found it difficult to identify the hydroxylated products of withanolides due to the similarity of masses and fragmentation pattern with other withanolides. Nevertheless, determination of the exact chemical structure of these products could shed some light into the role of WsCYP71B35 in the withanolide biosynthetic pathway.

Since biochemical assay products could not be confirmed, performing *in planta* studies would provide a better understanding of the gene function. To this end, VIGS and transient overexpression have been successfully adapted to study the *in planta* function of genes involved in the biosynthesis and regulation of withanloides in *W. somnifera.* Hence, *WsCYP71B35* was specifically silenced to determine its role in withanolides formation. It was found that the degree of *WsCYP71B35* silencing was comparable to the VIGS of other related genes of *W. somnifera* reported in earlier studies by other groups as well as our own previous studies (Singh et al. 2015, 2017; Agarwal et al. 2018; Shilpashree et al. 2022). Silencing of *WsCYP71B35* resulted in a significant reduction of withanolide A, while its overexpression resulted in a drastic increase in withanolide A along with a reduction in withanolide B content (Figure 5). This suggests that *WsCYP71B35* may play a direct role in withanolide A biosynthesis and have some indirect role in the regulation of other withanolide biosynthesis. Curiously, the hydroxylation of the 20^th^ carbon of withanolide B forms withanolide A. However, this is unlikely to be the activity of WsCYP71B35 as enzyme assay with withanolide B clearly showed products at a much different retention time (Figure 4). Also, the fact that this enzyme did not form any detectable product when incubated with phytosterols indicated unknown steps and precursor/s in the pathway between phytosterols and withanolide A. Besides, though not statistically significant, withaferin A also showed some modulation as *WsCYP71B35* silenced plants showed increased levels, and overexpressing plants showed reduced levels. Withaferin A shares a common skeletal structure with withanolide B but lacks the C-20 hydroxylation found in withanolide A. Based on our metabolite analysis, we speculate withanolide B may appear early in the withanolide biosynthetic pathway and branched to withanolide A and withaferin A separately, and that *WsCYP71B35* plays a role in withanolide A branch of this complex pathway (Figure S3).

CYP450s play important roles in plant defense through their involvement in phytoalexin biosynthesis, hormone metabolism and the biosynthesis of some other secondary metabolites (XU et al. 2015). When we assessed the role of *WsCYP71B35* in plant defense, it was found that modulation of expression of *WsCYP71B35* not only modulated withanolide levels in the plant but also modulated the expression of defense related *PR* genes, *PR1* and *PR3*. Further, it was observed that VIGS of *WsCYP71B35* led to significantly reduced tolerance of plants to *P. syringae* growth and in contrast the overexpression of *WsCYP71B35* resulted in increased tolerance to the same bacterium (Figure 7). It is interesting to note that these results are similar to our previous characterization of three CYPs from *W. sominifera* indicating that cytochrome P450 enzymes play a crucial role in defense in this plant (Shilpashree et al. 2022). Moreover, AtCYP76C2 and AtCYP71A12 from *Arabidopsis* which were found to be associated with defense mechanism against *Pseudomonas syringae* infection (Godiard et al. 1998; Kempthorne et al. 2021). Hence, it is clear that *WsCYP71B35* is involved in bacterial defense, however the exact mechanism of signaling needs further investigation. It is possible that the products (one or more withanolides) formed by this enzyme could be involved in providing tolerance through their antibacterial activity or in signaling. Similar to our results, it has been previously reported that overexpression of *Panax ginseng PgCYP76C9* conferred enhanced resistance to *P. syringae* in transgenic Arabidopsis (Balusamy et al. 2017). When we overexpressed *WsCYP71B35* in a transgenic *N. tabacum*, we observed an increased tolerance to *P. syringae* infections. This shows that *WsCYP71B35* has an independent role in plant defense. In conclusion, this study identified and characterized a novel CYP450 enzyme from *W. somnifera* and gave some insight to the biosynthetic pathway of withanolides. This will aid future work in the field of synthetic biology for the recombinant production of withanolides. This study also gives insight into biotechnological approaches for improving withanolide content by gene-editing. We also show the role of *WsCYP71B35* in plant defense.

In summary, this study has presented biochemical evidence for the utilization of certain withanolides as substrates by WsCYP71B35. Moreover, it has demonstrated the *in planta* role of WsCYP71B35 in both withanolides biosynthesis and defense against a model bacterial pathogen. Further elucidation of the enzymatic products through Nuclear Magnetic Resonance (NMR) analysis would offer valuable insights into the precise biochemical nature of WsCYP71B35. The characterized CYP450 could also be utilized in the metabolic engineering of *W. somnifera* cell cultures or the whole plant for the targeted enhancement of specific withanolides, and for the development of plants tolerant to bacterial pathogens.

## Author contributions

H.B.S, A.K.N., S.R.K., and V.B. performed the experiments. H.B.S, A.K.N and D.A.N analysed the data. D.A.N conceived and coordinated the research. H.B.S, A.K.N., and D.A.N wrote the manuscript.

## Data availability

All the data generated or analysed during the study are available from the corresponding author upon request.

## Acknowledgements

H.B.S and A.K.N are recipients of research fellowships from Indian Council of Medical Research (ICMR). The authors are thankful to the Director, CSIR-CIMAP, for the support throughout the study. This manuscript bears the institutional communication number CIMAP/PUB/ 2023/147. Authors declare no conflict of interest

