## Supplemental Information for "The cytochrome P450 enzyme WsCYP71B35 from *Withania somnifera* has a role in withanolides biosynthesis and defense against bacteria"

**Supporting Information:**

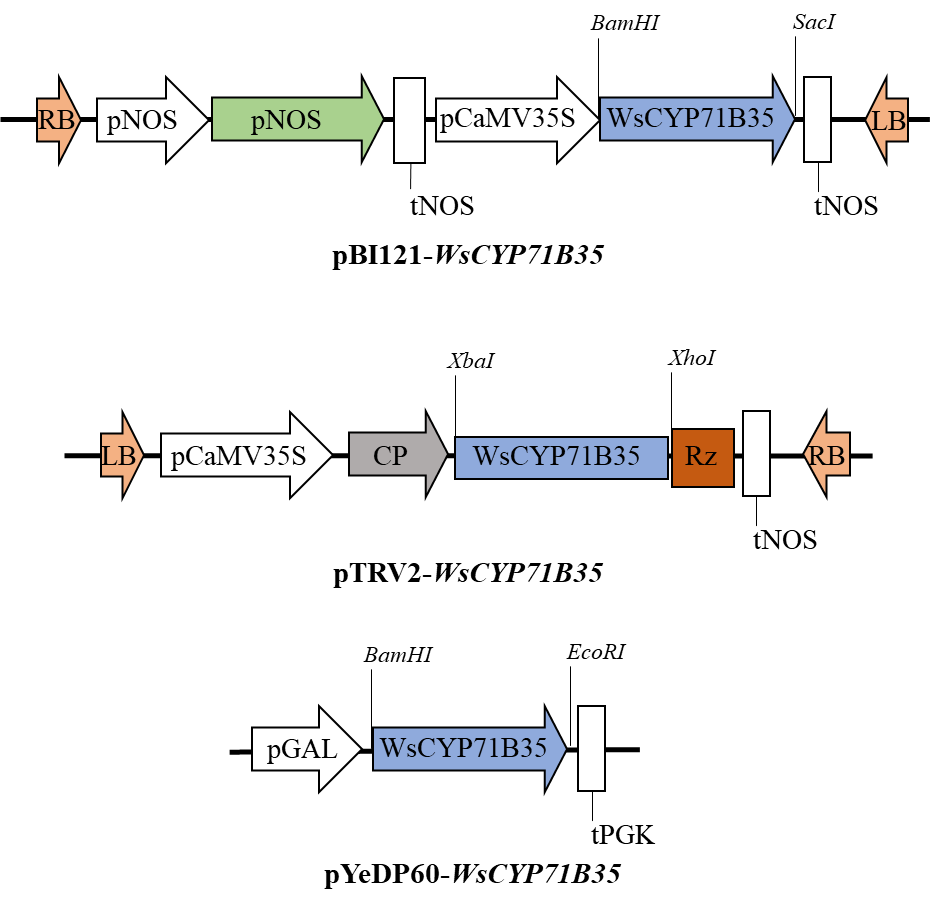

**Figure S1:** Pictorial representation of plant overexpression, VIGS and yeast expression constructs generated in this study.

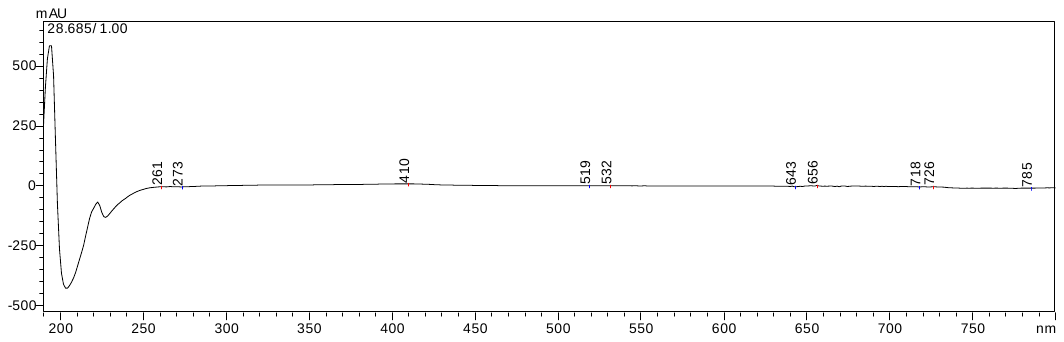

**A**

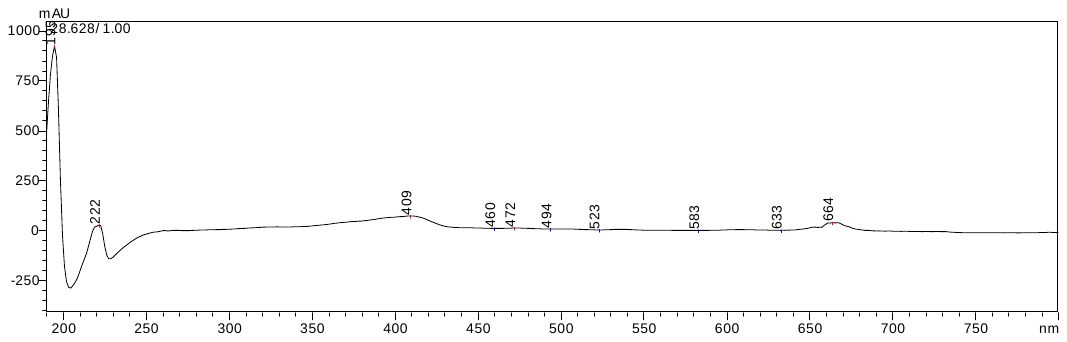

B

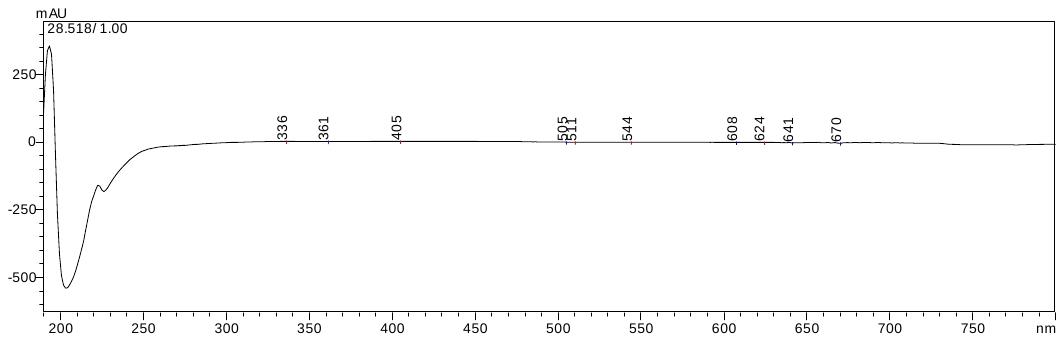

**C**

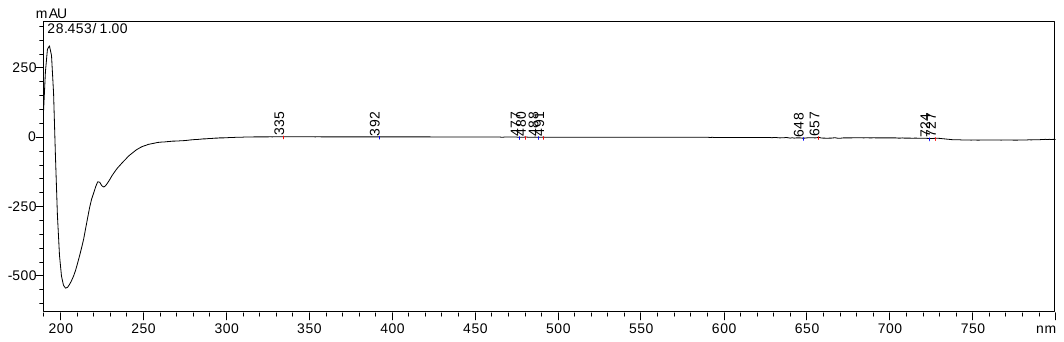

**D**

**Figure S2:** UV-Vis spectra of the product peaks formed by WsCYP71B35 when incubated with A) withaferin A, B) withanolide A, C) withanolide B and D) withanoside IV.

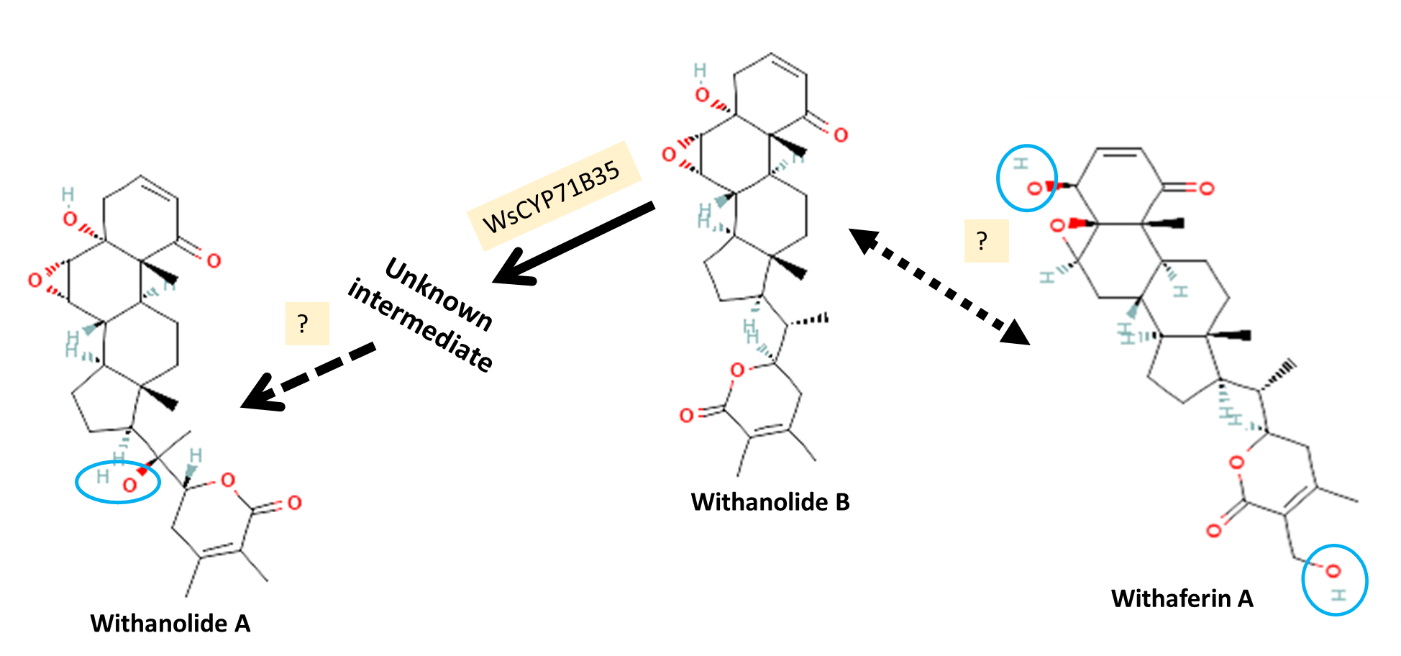

**Figure S3:** Possible role of WsCYP71B35 in withanolide biosynthetic pathway based on metabolite analysis of *WsCYP71B35* silenced and overexpressed samples. It is postulated that withanolide B gets converted to withanolide A and withaferin A through separate pathways. Key hydroxyl groups that differentiate withanolide A and withaferin A to withanolide B are marked with blue ovals.

**Table S1.** GenBank accession numbers for all sequences used in Figure 2:

| **Species** | **Gene Name** | **GenBank Accession No.** |
| --- | --- | --- |
| *A. thaliana* | AtCYP71A16 | AED94832.1 |
| *Ajuga reptans* | ArCYP71D443 | BAS30379.1 |
| *Ammi majus* | AmCYP71AJ1 | Q6QNI4.1 |
| *Arabidopsis thaliana* | AtCYP71 | NP_189318.1 |
| *Artemisia annua* | AaCYP71AV1 | BAM68808.1 |
| *C. intybus* | CiCYP71AV8 | E1B2Z9.1 |
| *Catharanthus roseus* | CrCYP71D12 | ACM92061.1 |
| *Cichorium intybus* | CiCYP71BL3 | G3GBK0.1 |
| *Hyoscyamus muticus* | HmCYP71 | ABS00393.1 |
| *Lactuca sativa* | LsCYP71 | AEI59780.1 |
| *Lotus japonicus* | LjCYP71D353 | AHB62239.1 |
| *Manihot esculenta* | MeCYP71E7 | Q6XQ14.1 |
| *Melia azedarach* | MaCYP71BQ5 | QDZ36307.1 |
| *Melia azedarach* | MaCYP71CD2 | QDZ36314.1 |
| *Mentha x gracilis* | MgCYP71D95 | Q6WKY9.1 |
| *Oryza sativa* | OsCYP71Z6 | A3A871.1 |
| *Populus trichocarpa* | PtCYP71B40v3 | AIU56748.1 |
| *Sorghum bicolor* | SbCYP71E1 | AAC39318.1 |
| *Vitis vinifera* | VvCYP71BE5 | BAT70338.1 |
| *Zingiber zerumbet* | ZzCYP71BA1 | BAJ39893.1 |

**Table S2:** List of oligonucleotide primers used in this study.

| **Primer** | **Sequence**  **5′ to 3′** | **Purpose** |
| --- | --- | --- |
| *WsCYP71B35*-RT_F | CTTTTCCCATTCGCTCGATTC | qRT- PCR |
| *WsCYP71B35*-RT_R | CTGAAGGTACATGGCAAGGTTGA | qRT- PCR |
| *WsPR1-*RT_F | GCTTCTCATCGACCCACATCTT | qRT- PCR |
| *WsPR1-*RT_R | GGAAAGCGGCGGCTAGA | qRT- PCR |
| *WsPR3-*RT_F | CCCCATGAATAGGGACCATCT | qRT- PCR |
| *WsPR3-*RT_R | GAGAAGTCTGAGCCAGAAAGGC | qRT- PCR |
| *WsCYP71B35-*VIGS F | TCTAGATCTTCTTCCTTCTCTACTGAAATGGAG | pTRV2 cloning |
| *WsCYP71B35-*VIGS R | CTCGAGCGACAACTCATCTTGACGAAT | pTRV2 cloning |
| *WsCYP71B35-*pYEDP60u F | GGGATCCATGGACTACAAAGACGATGACGACAAGG AGATTTATCAGTTGTTCAACTCTG | pYEDP60u cloning |
| *WsCYP71B35-*pYEDP60u R | GGGAATTCTCAAGAATAATAAGGAGTGGCAATC | pYEDP60u cloning |
| *WsCYP71B35-*PBI F | GGGATCCATGGAGATTTATCAGTTGTTCAACTCTG | pBI121 cloning |
| *WsCYP71B35-*PBI R | GGAGCTCTCAAGAATAATAAGGAGTGGCAATC | pBI121 cloning |

**Table S3:** Prediction of subcellular localization of WsCYP71B35 by different tools.

| **Tools** | **WsCYP71B35** |
| --- | --- |
| Predotar | ER |
| WoLF PSORT | chlo: 8, nucl: 2, plas: 2, cyto: 1, vacu: 1 |
| TargetP | Signal peptide, 0.5291 likelihood |
| iPSORT | Signal peptide present, Average Hydropathy (KYTJ820101), Value 1.935 |
